# *Sl*SWEET15 exports sucrose from phloem and seed coat in tomato to supply carbon for fruit and seed development

**DOI:** 10.1101/2020.11.09.374827

**Authors:** Han-Yu Ko, Li-Hsuan Ho, H. Ekkehard Neuhaus, Woei-Jiun Guo

## Abstract

Tomato, an important fruit crop worldwide, requires efficient sugar allocation for fruit development. However, molecular mechanisms for sugar import to fruits remain poorly understood. Expression of SWEET (Sugars Will Eventually be Exported Transporters) proteins is closely linked with hexose ratio in tomato fruits and may be involved in sugar allocation. Here, using quantitative PCR, we discovered that *SlSWEET15* was highly expressed in developing fruits compared to vegetative organs. Based on *in situ* hybridization and GUS fusion analyses, *Sl*SWEET15 proteins accumulated in vascular tissues and seed coats, major sites of sucrose unloading in fruits. Localizing *Sl*SWEET15-GFP to the plasma membrane supported its putative role in apoplasmic sucrose unloading. The sucrose transport activity of *Sl*SWEET15 was confirmed by complementary growth assays in a yeast mutant. Elimination of the *Sl*SWEET15 function by CRISPR/cas9 gene editing significantly decreased average sizes and weights of fruits, with severe defects in seed filling and embryo development. Together, we confirmed the role of *Sl*SWEET15 in mediating sucrose efflux from the releasing phloem to the fruit apoplasm and subsequent import into parenchyma cells during fruit development. Furthermore, *Sl*SWEET15-mediated sucrose efflux was also required for sucrose unloading from the seed coat to the developing embryo.

**One-sentence Summary:** *Sl*SWEET15, a specific sucrose uniporter in tomato, mediates apoplasmic sucrose unloading from releasing phloem cells and seed coat for carbon supply during fruit expansion and seed filling.

---

Tomato (*Solanum lycopersium*) is a key fruit crop worldwide with >$80B annual production value (FOASTAT, 2016). Development of tomato varieties with high yield and excellent quality have been primary targets for genetic improvement (Ruan et al., 2012; Wang et al., 2019). Photosynthetic assimilation supply is considered a major limiting factor for fruit development (Paul et al., 2018; Quinet et al., 2019). Up to 80% of fruit carbon is imported from source leaves (Hetherington et al., 1998), with sucrose the major form of carbon translocated to tomato fruits (Walker and Ho, 1977; Abbes et al., 2009; Osorio et al., 2014; Milne et al., 2018). Consequently, increased sucrose allocation to fruits is a potential strategy to increase yield and quality (Ruan et al., 2012; Osorio et al., 2014).

Tomato fruit development is typically divided into four stages: cell division, expansion, ripening and maturation (Pesaresi et al., 2014; Quinet et al., 2019). During early stage-cell division, from 0 to 14 days after anthesis (DAA), based on symplastic tracer and radiotracer studies, sucrose is mainly unloaded from sieve element-companion cell (SE-CC) lumens to surrounding vascular and storage parenchyma cells (PCs) via connecting plasmodesmata, with a small portion transported via an apoplasmic pathway (Ruan and Patrick, 1995; Patrick and Offler, 1996). However, during rapid expansion (14 to 40 DAA; Quinet et al., 2019), the young fruit switches to complete apoplastic sucrose unloading from the SE-CCs to PCs (Ruan and Patrick, 1995). Apoplastic sugar transport is indicated by a reduced abundance of plasmodesmata connections between SE-CCs and PCs and loss of mobility of symplasmic dyes in the area of release phloem of tomato pericarp cells (Johnson et al., 1988; Ruan and Patrick, 1995). Apoplasmic unloading of sugars continues for the remaining stages of tomato fruit ripening (Johnson et al., 1988). A feature of the expansion stage is high sugar accumulation, which requires extensive sucrose unloading (Walker and Ho, 1977; Damon et al., 1988; Quinet et al., 2019). This would require a plasma membrane-localized sugar transport mechanism to enable sucrose export from the release phloem to the fruit apoplasm (Lalonde et al., 2003; Osorio et al., 2014; Milne et al., 2018). Involvement of sugar carriers for apoplasmic unloading has been reported in several fruit crops (Braun et al., 2014; Milne et al., 2018), including cucumber (Hu et al., 2011), apple (Zhang et al., 2004) and grape (Wang et al., 2003).

However, the molecular carrier responsible for initial sucrose unloading from SE-CCs to fruit apoplasm has been elusive. Localization in the plasma membrane of SEs in tomato fruits implies that the sucrose transporter *Le*SUT2 may participate in sugar unloading in fruits (Barker et al., 2000; Hackel et al., 2006). Knockdown of *LeSUT2* expression caused a 20 to 40% reduction of both fruit sugar concentration and fruit size (Hackel et al., 2006), wheras overexpression of the pear sugar transporter *Pb*SUT2 in tomato enhanced sucrose concentrations and numbers of fruit produced (Wang et al., 2016). Nevertheless, based on the active transport properties of this symporter (Schulze et al., 2000; Carpaneto et al., 2010; Kuhn and Grof, 2010), *Le*SUT2 likely functions as a retrieval system to prevent sugar loss from SE-CCs, instead of exporting sucrose to the fruit apoplast (Hackel et al., 2006; Milne et al., 2018). Passive SWEET (<u>s</u>ugar <u>w</u>ill <u>e</u>ventually be <u>e</u>xported <u>t</u>ransporter) uniporters are likely candidate carriers responsible for initial apoplasmic phloem unloading in sink fruits (Osorio et al., 2014; Milne et al., 2018). Latter hypothesis is consistent with the passive transport nature of sucrose unloading in tomato fruit (Damon et al., 1988; Johnson et al., 103 1988).

The SWEET gene family has been identified in a wide variety of plants, including tomato (Chen et al., 2015; Feng et al., 2015). Based on their amino acid sequences, SWEET proteins are divided into four distinct clades (Chen, 2013; Chen et al., 2015). Clades I and II mainly transport glucose, clade III could transport sucrose and clade IV can transport fructose. SWEET was first identified as the central player that mediates sucrose efflux from mesophyll cells to the apoplast prior to phloem loading (Chen et al., 2012; Bezrutczyk et al., 2018; Gao et al., 2018). Furthermore, the passive uniporter feature of SWEET members provides an energy-efficient mechanism for unloading sugar in sink organs. In Arabidopsis, *At*SWEET11, 12 and 15 are localized on plasma membranes of maternal integument and filial endosperm cells to mediate a cascade of sugar unloading that supports embryo development (Chen et al., 2015). Furthermore, *Os*SWEET11 and 15 in rice (Yang et al., 2018), *Zm*SWEET4c in maize (Sosso et al., 2015), and *Gm*SWEET15 in soybean (Wang et al., 2019) also participated in apoplasmic sucrose unloading in developing seeds to support endosperm and embryo development. In tomato, based on transient silencing and genetic analyses, *Sl*SWEET1a also participated in sucrose unloading to sink leaves, as well as regulating the fructose/glucose ratio in ripening fruits (Shammai et al., 2018; Ho et al., 2019). Similarly, expression of pear *Pu*SWEET15 was closely linked with sucrose contents in pear fruit (Li et al., 2020). These findings led us to hypothesize that a *SlSWEET* member expressed in developing tomato fruits is involved in sucrose unloading for fruit development.

In this study, we determined that *SlSWEET15*, belonging to clade III, was highly expressed during the expansion stage of fruit development. Both *SlSWEET15* RNA and protein were localized in vascular tissues of most fruit tissues. Moreover, *Sl*SWEET15 protein was also present in seed coat and ripening fruits, implicating *Sl*SWEET15 in sugar transport throughout fruit development. Expression in yeast demonstrated that *Sl*SWEET15 probably functions as a sucrose-specific transporter on the plasma membrane. Knocking-out the *Sl*SWEET15 function, by CRISPR-Cas9-mediated gene editing, retarded fruit development and impaired seed filling. In summary, our findings indicate that the *Sl*SWEET15 facilitator had an essential role in sucrose unloading from releasing phloem to support fruit and seed development in tomato.

## RESULTS

### *SlSWEET15* was highly expressed in phloem cells of developing fruits

To identify which *SlSWEET* gene is critical for sugar unloading during fruit expansion, cDNA samples for expression profiling were prepared from immature green tomato fruits 14 days after anthesis (DAA). Based on preliminary quantitative reverse transcription (qRT)-PCR analysis, *SlSWEET15*, which belongs to clade III *SWEET* members, was strongly expressed during fruit expansion when compared to all other *SlSWEET* genes (Supplemental Fig. S1). In expression profiles of clade III *SlSWEETs* during fruit development (14 to 42 DAA), *SlSWEET15* had the highest expression in developing green fruits (14-21 DAA), but relatively low expression in mature ripening fruits (35-42 DAA, Fig. 1A). Expression of *SlSWEET15* was low in vegetative organs (e.g. roots and leaves; Fig. 1B) and only slightly in flower organs (Fig. 1B). Based on these results, we inferred that *SlSWEET15*a had a specific role during fruit development.

**Figure 1.**
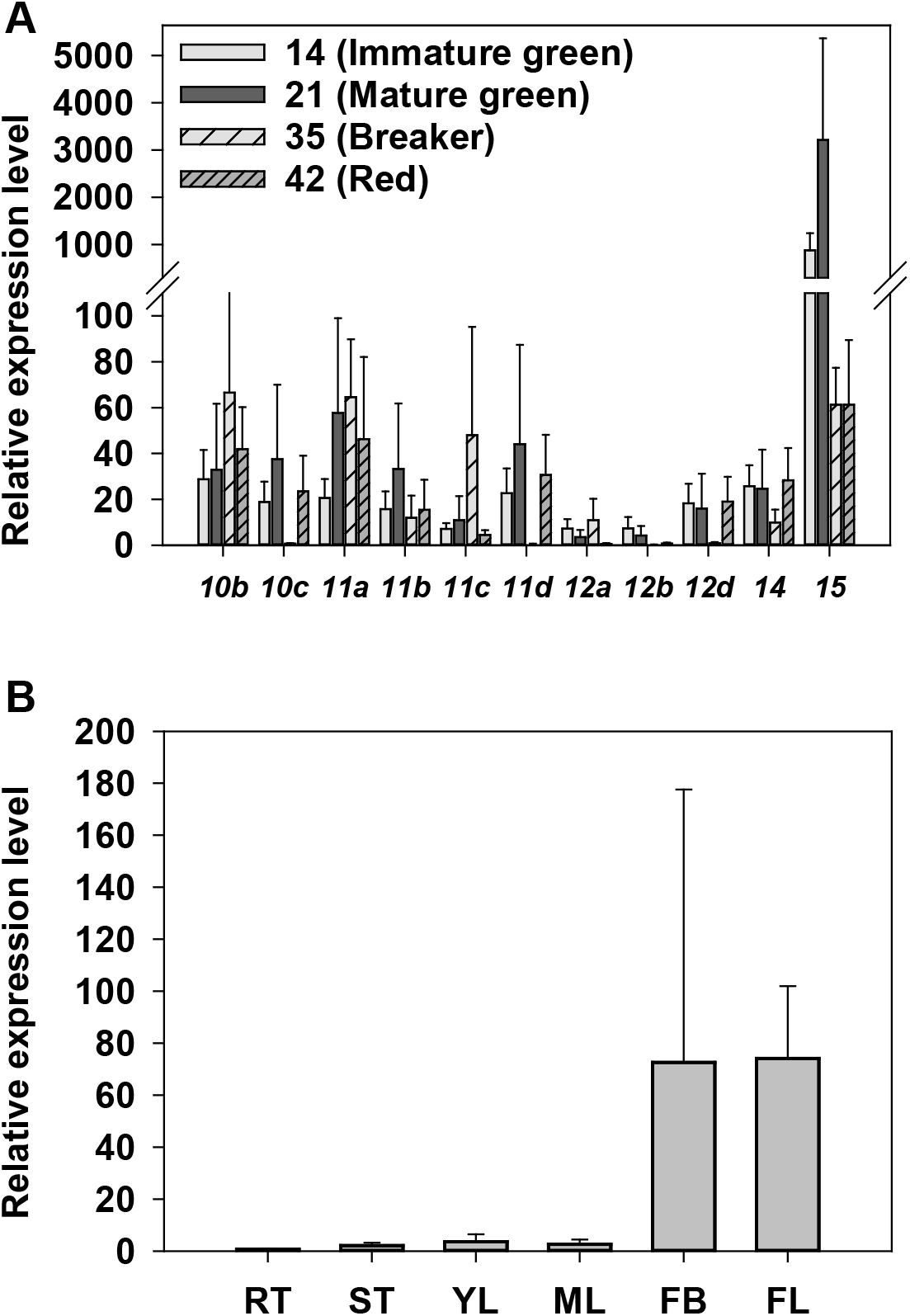
Expression of *SlSWEET15* in tomato organs. A, Expression of representative clade III *SlSWEET* genes in various stages of tomato fruits. B, Expression of *SlSWEET15* in non-fruit organs. Total RNA was isolated from fruits of 14, 21, 35, 42 DAA (A) or various organs and the derived cDNA was used for qRT-PCR with specific primers. The ordinate is the relative expression level, normalized to the internal control SlActin7. Results are mean ± SE from 3-7 independent biological repeats. DAA, day after anthesis. RT, root. ST, stem. YL, young leave. ML, mature leave. FB, flower bud. FL: flower.

To address where *SlSWEET15* was expressed, tissue-specific localization of *SlSWEET15* transcripts was examined in mature green fruits (21 DAF) by *in situ* hybridization with gene-specific probes. Compared to the background condition (sense probe), anti-sense signals were detected in all fruit cells (Fig. 2A). In particular, there were substantial signals in phloem cells and neighboring PCs in pericarp, columella, placenta, as well as seed coat (arrow heads in Fig. 2A; Supplemental Fig. S2A).

**Figure 2.**
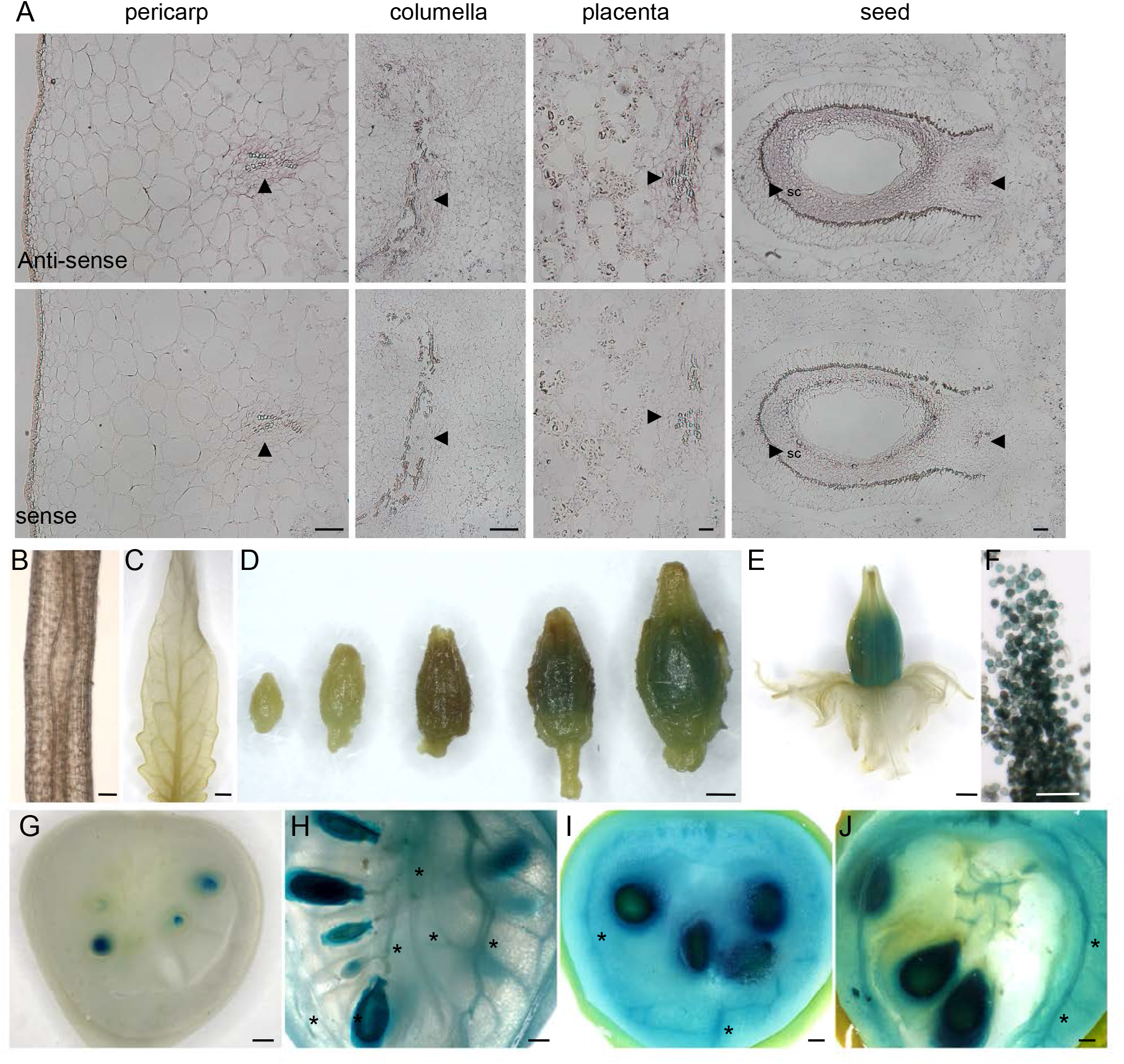
Organ-specific expression patterns of SlSWEET15 in tomato. A, Cell-specific localization of *SlSWEET15* transcripts analyzed by *in situ* hybridization. Cross-sections of tomato fruits (21 DAA) were hybridized with *SlSWEET15-*specific anti-sense (top) and sense probes (bottom). Arrowheads indicate locations of signals observed. B-J, Histochemical staining of GUS activities in transgenic tomato plants expressing SlSWEET15–GUS fusion proteins driven by *SlSWEET15* native promoter. B, mature roots. C, young leaflet. D, flower buds. E, mature flower. F, pollens. G-J, fruits of 14, 21, 35, 42 DAA. Asterisks indicate localization of signals in vascular tissues. Bars = 100 µm in A, B and F and 1 mm in C, D, E and G-J.

### *Sl*SWEET15 proteins accumulated in sugar unloading cells

For several SWEET members, there are varying ratios of gene expression and protein abundance exist (Abbes et al., 2009; Guo et al., 2014; Chen et al., 2015). To examine protein expression, we generated transgenic tomato plants expressing *Sl*SWEET15-GUS fusion proteins that were derived from the full *SlSWEET15* genomic DNA sequence under control of its native promoter. In T1 transgenic plants, *Sl*SWEET15-GUS fusion proteins were in very low abundance in roots, leaves or developing flower buds (Fig. 2B-D). In mature flowers, however, histochemical staining readily detected GUS fusion protein in pollen (Fig. 2D-F). In young developing fruits (14 to 21 DAA), *Sl*SWEET15-GUS protein was highly abundant in seed coats, with moderate amounts in vascular tissues of pericarp or placenta (asterisks in Fig. 2G, H; Supplemental Fig. S2B). During fruit maturation (35 to 42 DAA), in addition to seed coat and vascular tissues, GUS activity was also detected in all pericarp cells (Fig. 2I-J). Substantial accumulation of the GUS fusion protein in vascular tissues and seed coats strongly implicated the *Sl*SWEET15 transporter in sugar unloading in fruits and during seed development.

### Dual targeting *Sl*SWEET15 to the plasma membrane and the vacuolar membrane

To examine how *Sl*SWEET15 may participate in sugar transport, subcellular localization of C-terminal translational GFP-fusions of *SlSWEET15* (*Sl*SWEET15-GFP) was examined in Arabidopsis protoplasts. When co-transformed with the plasma membrane marker *At*PIP2A-RFP (Nelson et al., 2007), green fluorescence signals derived from *Sl*SWEET15-GFP fusion overlapped with red fluorescence derived from *At*PIP2A-RFP fusions (arrowhead in the top row of Fig. 3). When cells were lysed, red signals from the plasma membrane marker, *At*PIP2A-RFP, were greatly reduced (Supplemental Fig. S3). However, green fluorescence was still observed in a cell internal structure, likely to be the vacuolar membrane. To examine this possibility, the vacuolar membrane marker AtγTIP-RFP {Jauh, 1999 #555} was co-transformed with *Sl*SWEET15-GFP. Consistently, green fluorescence colocalized with red signals derived from the *At*γTIP-RFP fusion on the tonoplast, in either intact or lysed protoplasts (arrow-heads in the middle and bottom rows of Fig. 3), and lined the inside of chloroplast (asterisk in the middle row of Fig. 3). Latter results suggest that *Sl*SWEET15 proteins probably mediate sugar transport function on both, the plasma membrane and tonoplast, in tomato fruit cells.

**Figure 3.**
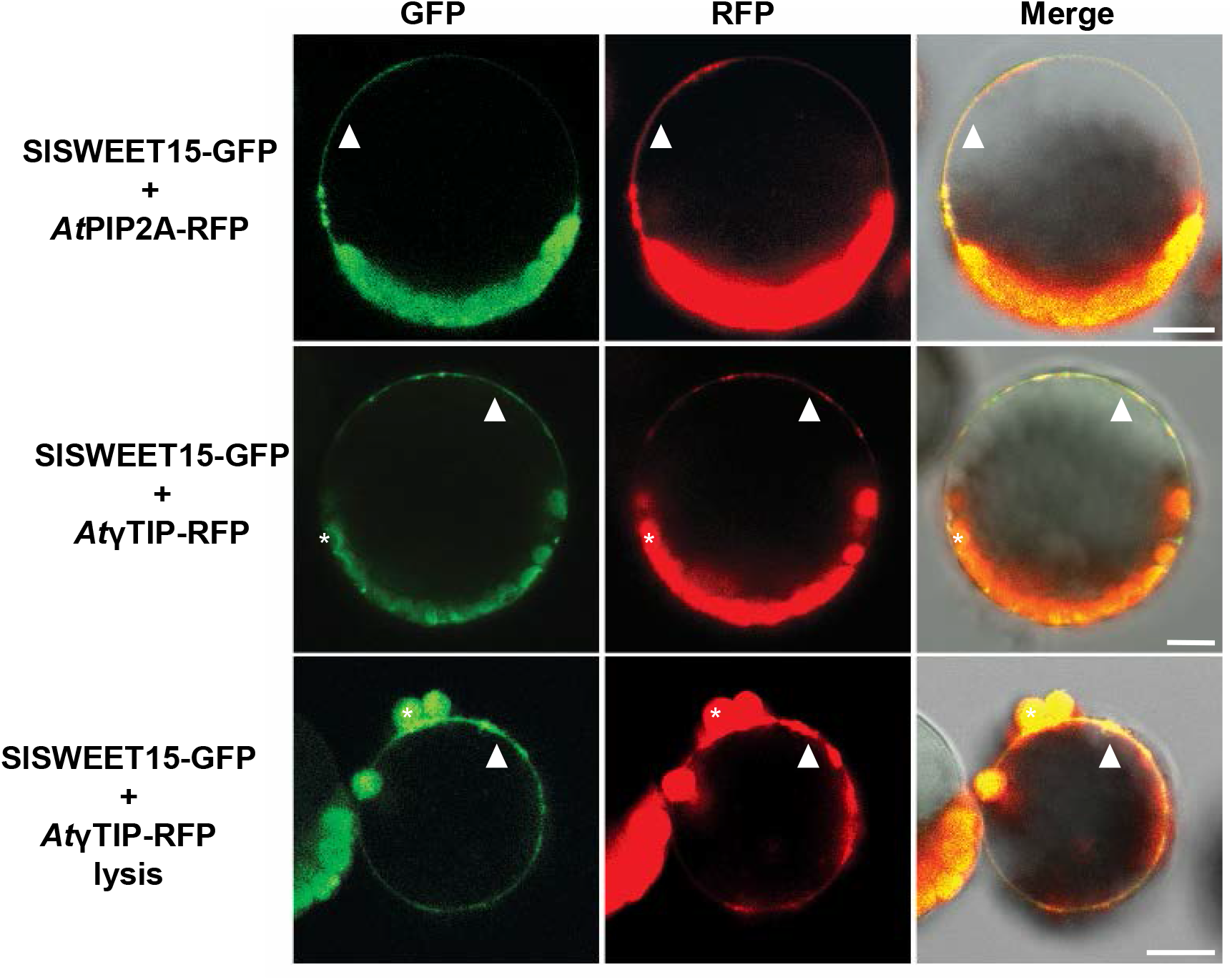
Dual localizations of SlSWEET15 in Arabidopsis protoplasts. Green and red fluorescence in Arabidopsis protoplast expressing SlSWEET15-GFP fusion proteins and *At*PIP2A-RFP or *At*γTIP-RFP fusions indicated localization to the plasma membrane and vacuolar membrane, respectively. Images of cells under lysis are also shown. Asterisk (*) and arrowheads indicate chloroplasts and overlapped signals, respectively. Bar = 10 µm.

### Transport activity of *Sl*SWEET15

When comparing amino acid sequence similarity, tomato *Sl*SWEET15 had high identity (up to 50%) to the Arabidopsis homolog *At*SWEET15 (Supplemental Fig. S4), a sucrose transporter (Chen et al., 2012). To examine transport activity, *SlSWEET15* was expressed in the bakers-yeast (*Saccharomyces cerevisiae*) mutant YSL2-1, which lacks all endogenous hexose transporters and the extracellular invertase. Accordingly, this mutant yeast was unable to grow on medium containing hexoses or sucrose (Chen et al., 2015). However, expression of the Arabidopsis sucrose transporter *At*SUC2 or yeast hexose transporter (HXT) restored growth on sucrose and hexose-containing medium, respectively (Fig. 4; Supplemental Fig. S5). *SlSWEET15* also complemented the growth deficiency of YSL2-1 on medium containing 2 or 5% sucrose (Fig. 4). Conversely, no yeast growth was present on glucose, fructose or galactose-containing medium (Supplemental Fig. S5), implying that *Sl*SWEET15 functioned as a sucrose-specific membrane transporter.

**Figure 4.**
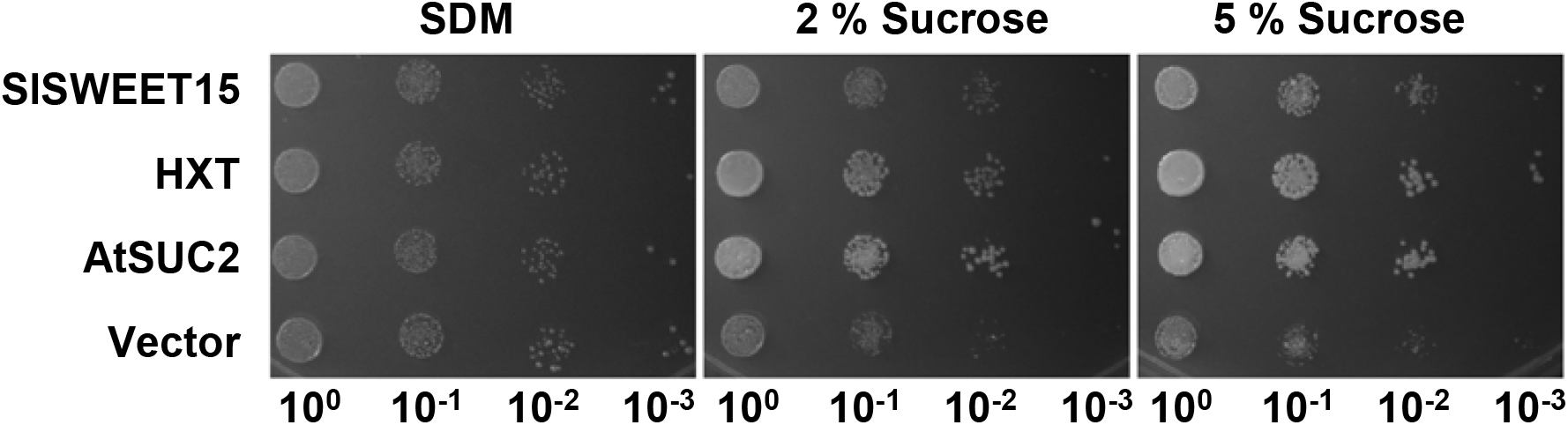
Sucrose transport activity of SlSWEET15 in yeast. Growth assay of YSL2-1 cells expressing SlSWEET15. Yeast cells expressing *SlSWEET15, HXT, At*SUC2 or an empty vector (vector) were serially diluted (10-fold) and cultured on solid media supplemented with maltose (SDM) or 2 to 5% sucrose, respectively. Images were captured after incubation at 30°C for 4 to 6 d.

### Establishment of *Sl*SWEET15 mutants using the CRISPR/Cas9 system

To provide genetic evidence of *Sl*SWEET15 function during fruit development, the CRISPR/Cas9 (clustered regularly interspaced short palindromic repeats and CRISPR-associated protein 9) gene-editing strategy was used to generate knockout mutants (Brooks et al., 2014). The plasmid was designed to produce two guide RNAs (T1 and T2-gRNAs), that would target +266 and +323 positions in exon 3 of *SlSWEET15*, with potential for a substantial deletion (red triangles in Fig. 5A; Brooks et al., 2014). To screen for editing mutations, we performed PCR on 15 individual T0 transgenic plants (Cas-15-1 to -19) using primers flanking both gRNA targets (Fig. 5A, P1 and P2). Lines 5 and 12 were identified as potentially having distinct deletions (Supplemental Fig. S6A). Detailed DNA sequencing of all PCR products confirmed that Cas-15-5, 15-12, 15-14 were bi-allelic knock-out mutants, whereas Cas-15-6, 15-7, 15-11 were heterozygous mutants (Fig. 5B). Mutations included deletions and point mutations (Fig. 5B). The Cas9 gene was detected in all mutant lines (the lower band in Supplemental Fig. S6A). Non-mutated transgenic plants that contained the transgene construct (denoted –V, e.g. Cas-15-1-V) or lacked the transgenes (denoted –WT, e.g. Cas-15-2-WT) were used as control plants for growth comparisons.

**Figure 5.**
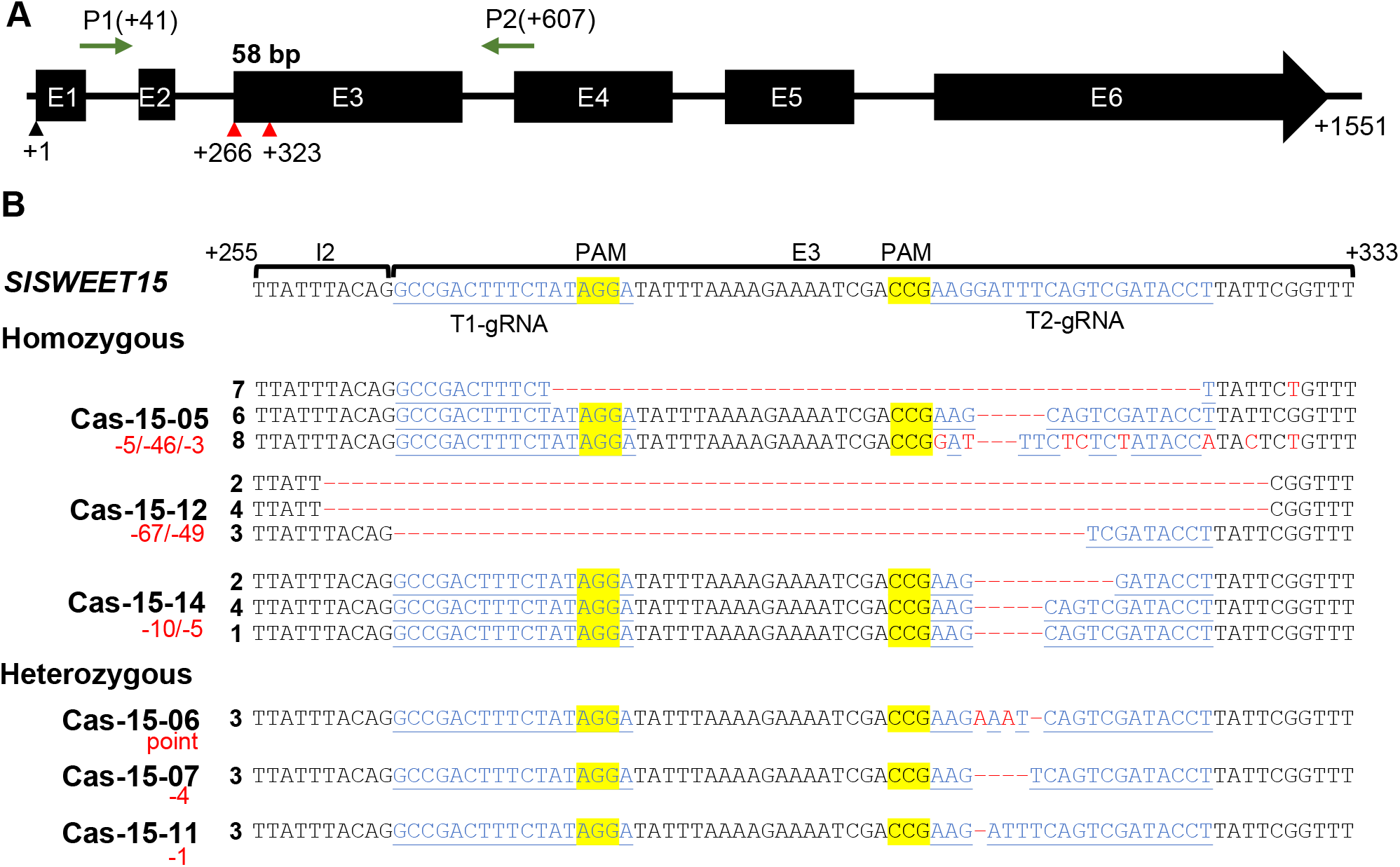
Genotypes of stable *slsweet15* mutant lines via CRISPR/Cas9 gene editing. A, Genomic structure of *SlSWEET15*. P1 and P2 indicated primers used to examine mutation types. Red triangles represented positions of two sgRNA sequences used in the binary vector. All numbers denoted the position of the first sequence relative to the start codon. B, Mutation types of *SlSWEET15* DNA in transgenic plants. Genomic DNA was isolated from mature leaves of T0 transgenic plants (Cas-15-x) and used to amplify gene fragments flanked with P1 and P2 primer in (A). Sequences of the resulting products from each line are shown. Sequences in blue and yellow represented target1/target2 (T1/T2) gRNA and PAM sequences, respectively. Red dash lines and font indicated deletion and point mutation, respectively. Numbers in red highlighted mutation types. E, exons. I, introns.

### Loss of *Sl*SWEET15 function inhibited fruit development

There were no consistent growth differences in plant size between T0 mutated transgenic tomato plants (Cas-15-5, 15-12, 15-14) and wild type plants (Supplemental Fig. S6B). However, mature red fruits derived from three independent homozygous knock-out lines were significantly smaller in size (polar and equatorial lengths, Fig. 6A) and fresh weight (∼40%, Fig. 6B). Interestingly, fruit size and weight were also reduced in heterozygous mutant plants (Fig. 6B). In addition, most seeds derived from homozygous knockout fruits had suppressed development (data not shown). The few seeds that did develop were smaller and flaky compared to those from non-mutated plants (WT/WT-V) (Fig. 6C). Consequently, seed weights of mutant plants were greatly decreased by 90% (Fig. 6D). The embryo and cotyledons had not developed in the mutant seeds (Cas-15-12) compared to non-mutated seeds (WT-V), where a well-developed embryo was enclosed by a thin layer of endosperm and seed coat (Fig. 6E). These T1 flaky seeds were unable to germinate even when supplied with sugars (data not shown). Unfortunately, no further phenotypes can be examined in T1 transgenic plants. Heterozygous mutant plants also had the same seed defects as those in homozygous plants (Fig. 6C), suggesting that *Sl*SWEET15 likely must form homotrimeric complex to be functional, as reported for a rice *Os*SWEET (Tao et al., 2015; Gao et al., 2018).

**Figure 6.**
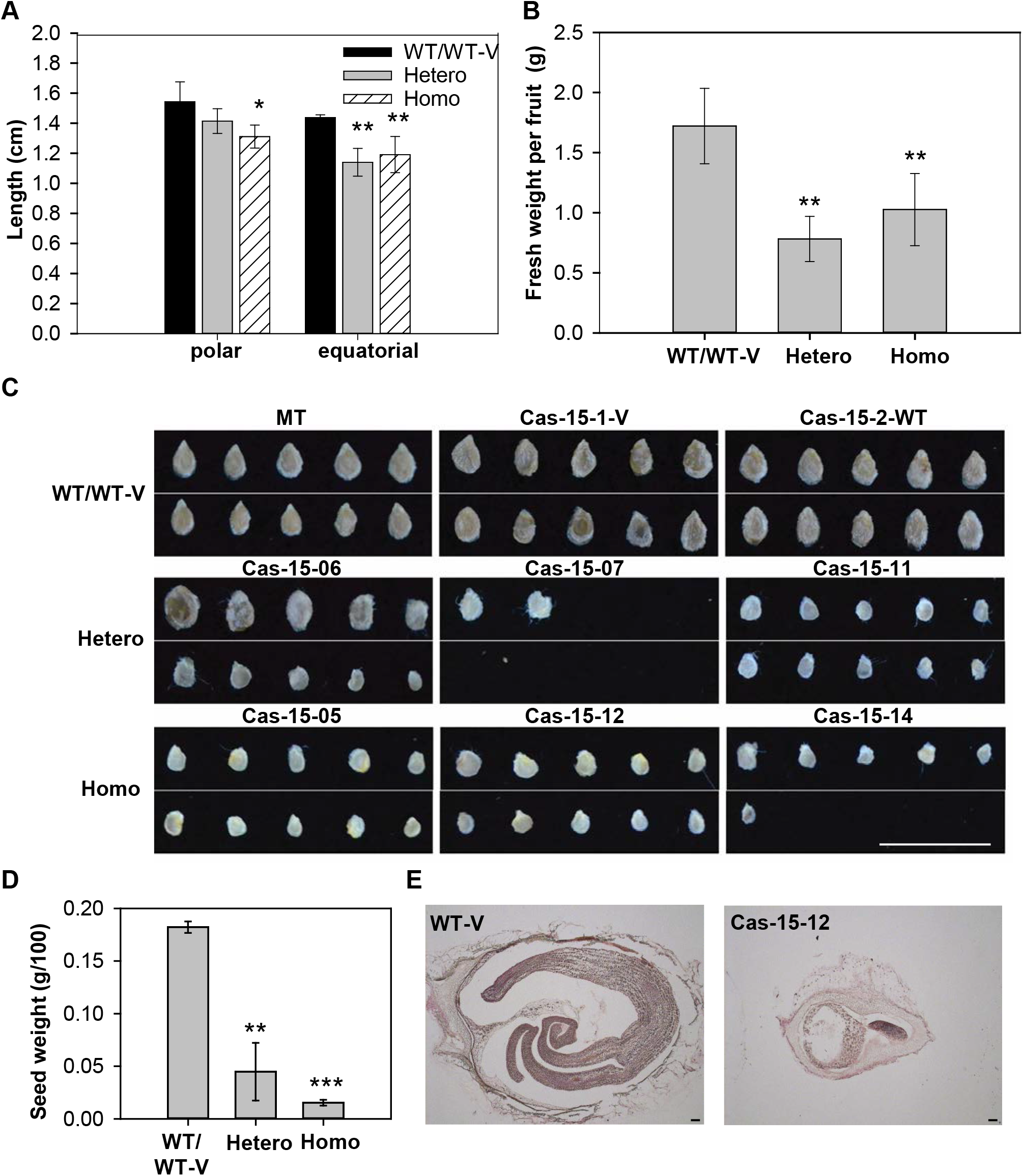
Fruit and seed development in slsweet15 knockout mutant plants. A and B, Fruit quality of T0 transgenic tomato plants. Mature red fruits were harvested and polar diameters, equatorial diameters (A) and fresh weights (B) were measured. C, Seeds of T0 transgenic tomato plants. Pictures are of representative seeds derived from wild type Micro-Tom (MT), transgenic plants without mutation (Cas-15-1-V and Cas-15-2-WT) and homozygous (Homo) or heterozygous (Hetero) mutants. Bar = 1 cm. D, Seed weight of T0 transgenic tomato plants. E, Longitudinal sections of wild-type and mutant seed. Bar = 100 μm. WT, WT-V indicated transgenic plants containing non-mutated genotype without or with the vector. Differences from WT (Student’s *t*-test): *(P<0.05), **(P<0.01) or ***(P<0.001) asterisks.

**Figure. 7.**
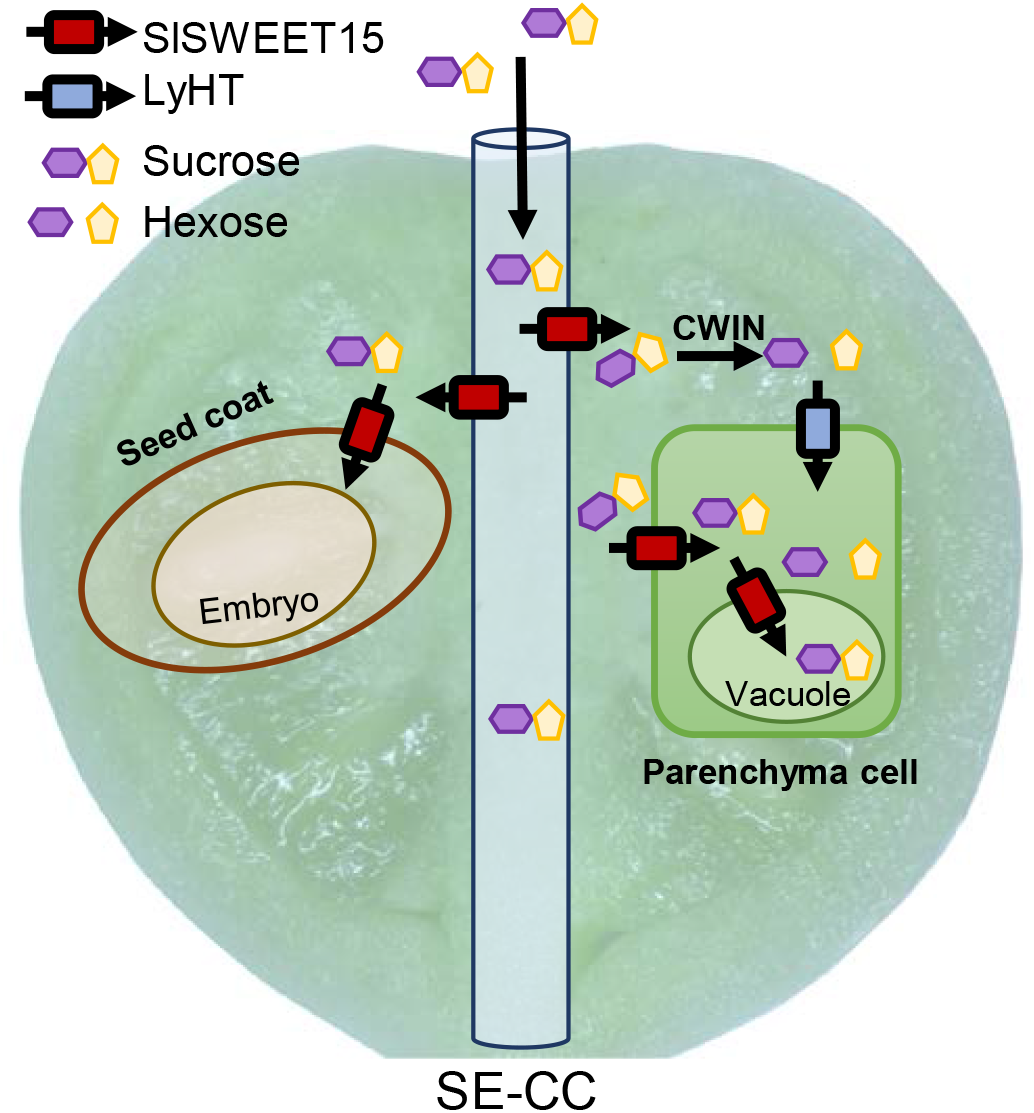
A functional model of SlSWEET15 during fruit and seed development. When sucrose is translocated from source leaves to developing tomato fruits, SlSWEET15 located on the plasma membrane of sieve-companion (SE-CC) complexes facilitated sucrose unloading to the fruit apoplasm. In addition, a portion of apoplasmic sucrose could be passively imported by the SlSWEET15 uniporter into neighboring parenchyma cells. Conversely, sucrose is also hydrolyzed by cell wall invertase (CWIN) and imported by active hexose transporter (HT). In seeds, SlSWEET15 also functions on the seed coat plasma membrane to mediate sucrose importation into an embryo for seed filling. During fruit maturation, SlSWEET15 was also located on the vacuolar membrane of pericarp cells and regulated intracellular sucrose dynamics.

## DISCUSSION

### *Sl*SWEET15 was involved in vascular sugar transport during fruit development

In tomato fruits, sugars are critical for fruit yield and quality (Pesaresi et al., 2014; Quinet et al., 2019). In particular, once reaching the rapid fruit growth stage, high sugar concentrations will quickly accumulate in pericarp or placenta cells, generating turgor pressure required for fruit expansion (Obiadalla-Ali et al., 2004; Quinet et al., 2019). During tomato fruit development, the cellular pathway of sucrose efflux from the releasing phloem SE-CCs switches from a symplasmic pathway to a complete apoplasmic pathway (Ruan and Patrick, 1995; Patrick, 1997). Accordingly, a plasma-membrane sucrose carrier in phloem cells is required, yet still elusive. In the current study, we concluded that *Sl*SWEET15 was the candidate sucrose carrier to mediate initial sucrose unloading for fruit expansion and seed filling in tomato.

Within all *SlSWEET* members, *SlSWEET15* transcripts were the sole isoform highly expressed in reproductive organs, in particular, in fruits that undergo expansion (Fig. 1; Supplemental Fig. S1). *In situ* hybridization analysis documented that during fruit expansion, the *SlSWEET15* transcripts accumulated in all releasing phloem units, namely the SE-CC complexes and surrounding PCs of major fruit tissues, e.g. pericarp, columella, and placenta (Fig 2A; Supplemental Fig. S2A). The pericarp is composed of layers of large, highly vacuolated parenchymatic cells (Czerednik et al., 2012), where sugars largely accumulate for cell expansion. The pericarp accounts for ∼50% of the fruit fresh weight at the expansion phase (Gillaspy et al., 1993; Obiadalla-Ali et al., 2004), making a substantial sugar influx mandatory for controlled fruit development. The vascular bundles of fruit columella represent the primary structure connecting the whole-plant vascular system and is the first location in fruits where sugars are unloaded from long-distance allocation (Gillaspy et al., 1993; Baxter et al., 2005). Sucrose and starch consistently accumulate in columella cells during fruit expansion (Baxter et al., 2005; Lemaire-Chamley et al., 2019). The placenta, interfacing maternal tissue, is clustered with vascular cells and parenchymatic tissues, from which the ovule primordia develops and the seeds are attached (Gillaspy et al., 1993; Brukhin et al., 2003). During fruit expansion, sucrose concentrations are generally higher in placentae than in the pericarp (Obiadalla-Ali et al., 2004). Thus, accumulation of *SlSWEET15* transcripts in phloem cells of sugar accumulating fruit tissues was closely associated with increased sugar import activity during the fruit expansion stage, which is almost twice as much as present during fruit maturation (Walker and Ho, 1977). In sum, our observations on gene expression, protein abundance and sugar accumulation implicated *Sl*SWEET15 in apoplasmic sugar transport in phloem cells during fruit expansion.

### *Sl*SWEET15 may mediate apoplasmic sugar unloading from phloem cells

The role of *Sl*SWEET15 in phloem was supported by specific accumulation of *Sl*SWEET15 proteins in vascular bundles of tomato fruits, based on a whole-gene *Sl*SWEET15-GUS translational fusion (Fig. 2B-J and Supplementary Fig. S2B). Additionally, co-localization of *Sl*SWEET15-GFP fusion proteins with a plasma membrane marker *At*PIP2-RFP (Fig. 3) revealed that *Sl*SWEET15 was located on the plasma membrane, the major site catalyzing apoplasmic sugar transport (Osorio et al., 2014; Milne et al., 2018). Localization of SlSWEET15 on the plasma membrane was consistent with subcellular localization of other SWEET15 homologs, e.g. Arabidopsis *At*SWEET15 (Chen et al., 2015) and pear *Pu*SWEET15 (Li et al., 2020), as well as other clade III SWEET members involved in sucrose loading in source leaves (Chen et al., 279 2010; Chen et al., 2012).

Most importantly, the constant accumulation of *Sl*SWEET15 proteins after 21 DAA in fruits was fully consistent with the transition timeframe of sucrose unloading from a symplasmic route in fruits at 13 to 14 DAA to a complete apoplasmic route at 23 to 25 DAA (Johnson et al., 1988; Ruan and Patrick, 1995; Patrick, 1997). Such apoplasmic sugar unloading from the phloem also persists throughout fruit maturation (Johnson et al., 1988). Based on accumulation of transcripts and proteins in phloem cells, as well as localization on the plasma membrane, we propose that *Sl*SWEET15 participates in apoplasmic sugar unloading required for development of tomato fruits. Furthermore, due to the additional tonoplast-localization of *Sl*SWEET15, perhaps *Sl*SWEET15 also participated in intracelluar sugar homeostasis in fruit cells, in particular during fruit maturation (Fig. 2I, 290 J).

### *Sl*SWEET15 may be a passive sucrose facilitator for sucrose accumulation

The sucrose-specific transport activity of *Sl*SWEET15, demonstrated by a yeast complemental assay (Fig. 4), supported its proposed role in sucrose unloading. Corresponding sucrose transport activity has been generally observed in almost all clade III SWEET transporters characterized, e.g. Arabidopsis *At*SWEET10-15 (Chen et al., 2012), cotton *Gh*SWEET10 (Cox et al., 2017), pear *Pu*SWEET15 (Li et al., 2020), Cassava *Me*SWEET10 (Cohn et al., 2014) and *Os*SWEET11 and 15 (Yang et al., 2018), *Zm*SWEET13 (Bezrutczyk et al., 2018). Furthermore, based on genetic evidence, sucrose contents in pear fruits were closed linked with expression levels of *Pu*SWEET15 (Li et al., 2020). These results highlighted a conserved sucrose transport feature of clade III SWEET transporters in both seed and fruit crops. Based on high identities (up to 48%) to its close Arabidopsis ortholog (*At*SWEET12, Km ∼70 mM, Supplementary Fig. S4B; Chen et al., 2012), *Sl*SWEET15 transporter probably also operated as a passive facilitator with low affinity, consistent with most characterized SWEET facilitators expressed in sink organs (Guo et al., 2014; Lin et al.,2014; Sosso et al., 2015; Ho et al., 2019). This passive transport fits well with the energy-independent transport nature of sucrose unloading in fruit pericarps that is not inhibited by metabolic inhibitors (e.g. PCMBS and CCCP; Damon et al., 1988; Brown et al., 1997; Obiadalla-Ali et al., 2004) and has low proton ATPase activities (Johnson et al., 1988). Moreover, high sucrose concentrations in SE cells, predicted to up to 500 mM and the steep concentration gradients (∼10 fold) of sucrose across the plasma membrane (Damon et al., 1988; Ruan and Patrick, 1995; Patrick, 1997), would favor a low affinity facilitator, like *Sl*SWEET15. Together with its localization at the plasma membrane of phloem cells, the passive sucrose-specific transport feature prompted us to conclude that *Sl*SWEET15 facilitated an energy-efficient sucrose-specific efflux from the plasma membrane of SE-CCs to fruit apoplasm to support fruit expansion.

Once sucrose is exported to the fruit apoplasm, it has been estimated that 70% of apoplasmic sucrose is hydrolyzed by cell wall inverses as hexoses, which should be quickly taken up into surrounding parenchyma cells by a hexose transporter in order to maintain a substantial sucrose concentration gradient for continuous unloading (Ruan and Patrick, 1995; Brown et al., 1997). Nevertheless, based on uptake of radioactive sucrose in pericarp cells, a small but significant portion of sucrose can be directly imported intact into fruit cytosols (Damon et al., 1988; Johnson et al., 1988; N’tchobo et al., 1999). The non-saturated and PCMBS-insensitive uptake feature of sucrose into fruit pericarp cells is consistent with an energy-independent carrier-mediated sucrose uptake, such as *Sl*SWEET15 (Damon et al., 1988; Johnson et al., 1988; Brown et al., 1997). Thus, it is likely that plasma-membrane and tonoplast localized *Sl*SWEET15 also facilitated sucrose import from the apoplasm to the cytosol and vacuole of storage parenchyma for continuous intra- or inter-cellular sugar allocation. Conversely, once sucrose is uploaded into fruit cells, most sugars stored in fruit vacuoles are glucose or fructose (Gillaspy et al., 1993; Milne et al., 2018; Vu DP et al., 2020). Notwithstanding, these stored hexoses could be exported to the cytosol and converted to sucrose, catalyzed by sucrose phosphate synthese (SPS) or Susy (N’tchobo et al., 1999). Changes in cellular sugar formats are ongoing, even during fruit maturation (N’tchobo et al., 1999). In this case, an energy-dependent sucrose uniporter, like *Sl*SWEET15, could provide an energy-dependent strategy to regulate sucrose homeostasis in fruit cells.

### *Sl*SWEET15 may function in sucrose unloading from the seed coat

Pronounced accumulation of *Sl*SWEET15 transcripts and proteins in seed coats and funiculus vascular cells implied that *Sl*SWEET15 was also involved in sucrose exchange in tomato seeds, which have strong nutrient requirements and are enriched with transporter proteins (Pattison et al., 2015). Requirement of a *Sl*SWEET15 transporter was consistent with a mandatory apoplasmic transport step between maternal (seed coat)-filial (endosperm or embryo) interface in tomato fruits (Ruan et al., 2012). Based on phloem-mobile fluorescent tracers or proteins, sucrose is apparently mostly unloaded from funiculus SEs symplasmically to seed coats (Patrick and Offler, 2001; Zhang et al., 2007), which develop from the ovule integument (Quinet et al., 2019). These integument cells enclose the embryo and are the major site for nutrient release in most developing dicot seeds, as reported in Arabidopsis (Stadler et al., 2005) and legume seeds (Wang et al., 1995). Yet, there is a symplasmic disconnection between outer and inner integuments in Arabidopsis (Werner et al., 2011), or between seed coat parenchyma cells and filial storage sites in legume seeds (Wang et al., 1995). A similar apoplasmic barrier was reported in monocot wheat and rice grains (Oparka and Gates, 1981; Wang and Fisher, 1995). In this scenario, a plasma membrane carrier, such as *Sl*SWEET15, would be required to mediate sucrose efflux to seed apoplasm. The proton-independent transport feature of the SWEET family is consistent with facilitated diffusion of sucrose efflux from seed coats (Zhang et al., 2007; Milne et al., 2018), as demonstrated in pea (De Jong et al., 1996; Zhou et al., 2007) and wheat (Wang and Fisher, 1995). The low affinity transport feature of SWEET proteins can also be physiologically favored, due to a substantial transmembrane sucrose concentration gradient (up to 50 mM) in maternal releasing cells in wheat grains and legume seeds (Fisher and Wang, 1995; Zhang et al., 2007). Based on these results, we inferred that *Sl*SWEET15 probably facilitated sucrose unloading from funiculus phloem cells and sucrose efflux from the seed coat cells for seed filling.

### *Sl*SWEET15–mediated sucrose export was required for fruit and seed development

Genetic evidence from CRISPR/Cas9 knock-out (KO) tomato mutants confirmed a physiological role of *Sl*SWEET15 in fruit development and seed filling (Fig. 5). A lack of *Sl*SWEET15 transport significantly reduced fruit growth and yield in Micro-Tom tomato (Fig. 6A, B), probably due an inadequate sucrose supply from the releasing phloem. Similarly, seeds were mostly aborted or flaky and lacked embryo development (Fig. 6C, D, E). These phenotypes were consistent with the role of SWEET15 analogs in non-fruit plants. In Arabidopsis seeds, *At*SWEET15, together with two clade III SWEET, *At*SWEET12 and 11, supported a cascade of sucrose transport from the outer and inner integuments to facilitate sucrose exchange from endosperm to embryo (Chen et al., 2015). In rice seeds, *Os*SWEET15 collaborated with *Os*SWEET11 to mediate sucrose unloading/export from vascular parenchyma into the apoplsmic space, enabling allocation and also export from the nucellar epidermis/eleurone interface to support seed filling (Yang et al., 2018). Defects in SWEET-mediated sucrose transport caused wrinkled and undeveloped seeds (Chen et al., 2015; Yang et al., 2018). Collectively, these studies supported a conserved function of clade III SWEET in sucrose unloading for seed development. In fleshy fruit crops, the same SWEET15 transport system to participate in sucrose unloading in both seed and fruit cells may reflect a close association between seed and fruit development, where signals derived from developing seeds control the rate of cell division in surrounding fruit tissues (Gustafson, 1939; Gillaspy et al., 1993)

## MATERIALS AND METHODS

### Plant and growth conditions

Tomato (*Solanum lycopersicum*) Micro-tom was used in this study. Tomato seeds were sterilized using a bleach solution (30% CLOROX and 0.1% Triton X-100) for 8 min and then washed twice with sterilized water. Tomato seeds were germinated in water for 2 to 3 d and transferred to soil mixture directly or to 1/2 MS liquid media (0.215% MURASHIGE & SKOOG MEDIUM, 0.1% MES, and 1.5% Agar) for hydroponics cultivation. All plants were grown in a controlled chamber (25°C, 16/8 h light/dark, with ∼100 μmol m^−2^ s^−1^ illumination). To analyze gene expression, various organs, including roots, stems, young leaves (<2 cm long) and mature leaves (>4 cm, terminal leaflet) were collected from 4-week-old tomato plants grown hydroponically. Flower buds (developing green buds), flowers (1DAA, 1 d after anthesis), fruits of 14 (immature green), 21 (mature green), 35 (breaker) and 42 (red) DAA were collected from 5-6 week old plants. All organs were stored at -80°C before analysis.

### RNA extraction

Fruit RNA transcripts were isolated according to the CTAB ((1-Hexadecyl)trimethyl-ammonium bromide) extraction method (Zhang et al., 2013). The extraction buffer contained 3% CTAB, 1.4 M NaCl, 20 mM EDTA, 100 mM Tris-HCl, 2% PVP40, and 2% β-Mercaptoethanol (pH 8). In short, samples were ground into powder and mixed with pre-heated CTAB extraction buffer and incubated at 65°C for 30 min. After centrifugation (8000 x *g* for 15 min), the supernatant was transferred to a new tube and mixed with an equal volume of chloroform:isoamylalcohol (24:1, v/v). The mixture was centrifuged (12000 x *g* for 30 min) and the supernatant was transferred and mixed with 1/3 volume of 10 M LiCl. The reaction was incubated at -20°C overnight. Pellets were collected by centrifugation, washed twice with 200 µL 4M LiCl, and suspended in 180 µL of 10 mM Tris-HCl (pH 7.5) and 20 µL of 3M potassium acetate (pH 5.5). These mixtures were kept on ice for 30 min and then centrifuged. The supernatant was transferred and mixed with the 2.5 volume of pre-cold isopropyl alcohol and stored at -70°C for 3 h. The RNA pellets were collected by centrifugation and washed with 75% ethanol and then dissolved in 20 µL DEPC-water.

RNA samples from other organs (except for fruits) were isolated using TRIzol reagent as instructed (Ambion® from Life Technologies). In short, samples were ground into powders, mixed with 500 µL TRIzol reagent and centrifuged. The mixtures were then transferred, mixed with 200 µL pre-cold chloroform:isoamyl alcohol (24:1, v/v) and centrifuged. The supernatant was added to 0.5 volume of 99% alcohol and resulting whole mixtures were transferred to an RNA spin column and processed as instructed (GeneMark, http://www.genemarkbio.com/). The RNA samples were suspended in 25 µL nuclease-free water and stored at -80°C until analyzed.

### Reverse transcription-PCR analysis

Total RNA transcripts were reverse-transcribed and gene-specific primers for 30 *SlSWEET* genes were used for real-time quantitative PCR (qRT-PCR), as described (Ho et al., 2019). The reference gene *SlActin7* was used to determine relative expression.

### *In situ* hybridization

To prepare the probe, partial *SlSWEET15* coding sequences of 246 bp were amplified with specific primers (RNAi-15-F and RNAi-15-R; Supplemental Table S1) and cloned into the vector pGM-T (Genomics). Digoxigenin-labeled sense and antisense RNA probes were synthesized following manufacturer’s instructions (Roche Applied Science). Mature green fruits (21 DAA) were sliced and fixed in pH 7.0 PFA solution (4% paraformaldehyde, 35 mM sodium hydroxide, 0.1% tween 20 and 0.1% triton X-100 in 250 mL PBS) for 16 h at 4°C. Samples were then dehydrated through an ethanol series and embedded into molten wax (Leica). Thick (10 μm) sections were cut on a MICROM 315R microtome (Thermo Scientific). Hybridization and immunological detection of signals with alkaline phosphatase were done as described (Lin et al., 2014).

### Expression of GUS fusions

The *SlSWEET15* (Solyc09g074530) promoter (2000 bp upstream to ATG) and genomic opening reading frame, including all introns (1348 bp after ATG) were amplified from genomic DNA with specific primers (SWT15-promoter-F and SWT15-promoter-R for promoter and SWT15-g-F and SWT15-g-R for open reading frame; Supplemental Table S1). The *Sl*SWEET15 promoter fragments were purified and cloned into the binary vector pUTKan by *SacI* and *SacII* sites (pUTKan-P_*SWEET15*_). The *Sl*SWEET15 genomic ORF was then cloned into pUTKan-P_*SWEET15*_ via SacII and BamHI sites. Tomato plants were transformed with the resulting pUTKan-P_*SWEET15*_::gSlSWEET15 binary vector in the Transgenic Plant Core Lab in Academia Sinica (http://transplant.sinica.edu.tw/en/-aboutus/intro/index3.htm) and three positive T0 transgenic tomato plants were obtained. Mature fruits of two T0 transgenic plants and various organs from heterozygous soil-grown T1 plants were collected and histochemically stained for 16 h, as described (Ho et al., 2019).

### Confocal microscopy for GFP fusions

To observe subcellular localization in yeast, the *SlSWEET15* cDNA fragment without the stop codon was amplified with a specific primer (5’UTR-SlSWT15 and attb-dTGA-R SWT15, Supplemental Table S1), then cloned into the pDONR221. The *Sl*SWEET15 cDNA was then transferred from the pDONR221 clone into p2GWF7 (Karimi et al., 2007). Arabidopsis protoplasts were isolated and transfected with resultant vector p2GWF7-SWT15, as described (Wu et al., 2009). To localize the position of inner membranes, plasma membrane marker *At*PIP2A:RFP fusions (Nelson et al., 2007), or the vacuolar membrane protein *At*rTIP:RFP fusion (Jauh et al., 1999) were also expressed with *Sl*SWEET15-GFP in protoplasts. After 20 to 34 h of transformation, fluorescence imaging of protoplasts was done on a Carl Zeiss LSM780 confocal microscope (Instrument Development Center, NCKU). The GFP fluorescence was visualized by excitation with at 488 nm and emission between 500 and 545 nm, whereas RFP fluorescence was visualized by excitation with at 561 nm and emission between 566 and 585 nm.

### Yeast complementation Assays

To express SlSWEET15 in yeast, cDNA sequence (861 bp) was amplified using Phusion polymerase (New England Biolabs) with gene-specific primers (attb-SlSWT15-F and SlSWT15-F). The cDNA was first cloned into the pDONR221-f1 vector using BP cloning and subsequently transferred to the pDRf1-GW vector using LR Gateway technology (Grefen et al. 2010). The yeast strain YSL2-1 was transformed with the resulting constructs (pDRf1-GW-SlSWT15) using the lithium acetate (LiAC) method (Chen et al. 2015). Transformants were selected and spotted on synthetic deficient media supplemented with or without various concentrations of sugars as described previously (Ho et al. 2019). Sequences of primers are provided in Table S1.

### Creating a *Sl*SWEET15 mutant line using CRISPR/Cas9

To create fragment deletion of *SlSWEET15*, two targeted sequences (T1 and T2), from positions +266 and +323 downstream of the translation start site (ATG), were chosen according to a website (https://crispr.cos.uni-heidelberg.de/; Fig. 5A). Targeted sequences were synthesized and the whole guide RNA scaffold including T1 and T2 sequences were amplified with specific primers (15-F0 and 15-R0) using the module vector pCBC-DT1T2 as a template. Resulting RNA scaffold products were used in an overlap PCR reaction with specific primers (SlSWT15-DT1-BsF and SlSWT15-DT2-BsR; Brooks et al., 2014). Thereafter, resulting PCR products containing the pCBC-DT1T2 *SlSWEET15-*specific cassette were digested with BsaI and inserted into the binary vector pKSE401 (Supplemental Figure S6Al Brooks et al., 2014), which was then introduced into an Agrobacterium strain and transformed into Micro-Tom tomato plant by the Transgenic Plant Core Lab in Academia Sinica (http://transplant.sinica.edu.tw/en/aboutus/intro/index3.htm). Nineteen positive T0 transgenic tomato plants were regenerated, transferred to soils and grown and fruits and seeds were collected. All primer sequences are shown (Supplemental Table S1).

### Genomic DNA extraction and PCR analysis

A mature leaf was collected from each T0 transgenic tomato plant and stored at -80°C pending analysis. Leaf samples were placed in liquid nitrogen, ground into powder and mixed with 500 µL CTAB extraction buffer (3% CTAB, 1.4 M sodium chloride, 2% PVP40, 20 mM pH8 EDTA and 100 mM pH8 Tris-HCl). Mixtures were incubated at 55°C for 15 min and centrifuged at 12000 x *g* for 5 min. The supernatant was transferred to new tubes, 250 µL chloroform:isoamyl alcohol (24:1, v/v) was added and the solution was vortexed and then centrifuged at 13000 x *g* for 1 min. The upper supernatant was removed and placed in 37.5 µL of 10 M ammonium acetate and 500 µL of pre-cold 99% alcohol and kept at -20 °C for 2 to 3 h, then centrifuged at 13000 x *g* for 1 min. Resulting pellets were washed twice with 70% alcohol, incubated at 60 °C for 5 min and finally re-suspended in 20 µL of nuclease-free water. To confirm mutation types in transgenic tomato plants, partial *SlSWEET15* fragments were amplified from genomic DNA with specific primers (P1 and P2) and cloned into the vector pGM-T, as instructed (Genomics, Taiwan). For each line, three to six derived clones were sequenced. To examine the transformation event, the gene sequence of Cas9 gene was also amplified with specific primers (Cas9-F and Cas9-R). All primer sequences are listed (Supplemental Table S1).

## ACKNOWLEDGEMENTS

This work was financially supported by grants from the Ministry of Science and 508 Technology, Taiwan (MOST 105-2628-B-006-001-MY3; MOST 108-2314-B-006-077-MY3) to W.J.G. Work in the lab of H.E.N was supported by the Deutsche Akademische Austauschdienst (DAAD, project PPP Germany 106-2911-I-006-506). We thank Dr. Peter Goldsbrough in Purdue University (IN, USA) for constructive comments on the manuscript.

## SUPPLEMENTAL MATERIALS

**Supplemental Table S1. The list of primers used in this study**.

**Supplemental Figure S1. Expression of *SlSWEETs* in developing tomato fruits**.

**Supplemental Figure S2. Expression pattern of *Sl*SWEET15 in developing tomato fruits**.

**Supplemental Figure S3. Subcellular localization of *Sl*SWEET15 in Arabidopsis protoplasts after lysis**.

**Supplemental Figure S4. Phylogenetic comparison of type III *SWEET* genes**.

**Supplemental Figure S5. Transport activities of *Sl*SWEET15 to hexoses in yeast**.

**Supplemental Figure S6. Identification of Cas9-mediated mutant plants**.

